# Cultivation and genomic characterization of the first representative of the globally distributed marine UBA868 group

**DOI:** 10.64898/2026.04.01.715867

**Authors:** Meora Rajeev, Yeonjung Lim, Mirae Kim, Dongkyun Kim, Ilnam Kang, Jang-Cheon Cho

## Abstract

Members of the UBA868 group within the order *Arenicellales* are globally distributed marine *Gammaproteobacteria* predicted to participate in sulfur and carbon cycling, yet their physiology and ecological roles remain unknown due to the absence of cultured representatives. Here, we report the isolation and characterization of the first heterotrophic representative of the previously uncultured UBA868 group. Using dilution-to-extinction cultivation, we obtained four isolates from the Yellow Sea whose high-quality genomes represent a single UBA868 species. One strain, IMCC57338, maintained in axenic culture, exhibited small coccoid morphology and slow growth (doubling time ∼2.9 days), consistent with an oligotrophic lifestyle. Genome analysis revealed a predominantly aerobic chemoorganoheterotrophic lifestyle with a streamlined central carbon metabolism, including a complete glyoxylate shunt and limited carbohydrate utilization capacity, suggesting adaptation to low-nutrient conditions. The genome also encodes pathways for methylated amine oxidation coupled to formaldehyde assimilation via the serine cycle, indicating a capacity for methylotrophy. Genes encoding sulfur oxidation (Sox) and reverse dissimilatory sulfite reductase (rDsr) pathways further suggest a capacity for sulfur-based chemolithoheterotrophy. Global metagenomic and metatranscriptomic read recruitment showed that the species represented by IMCC57338 is widely distributed across ocean basins and pelagic depth layers, with higher abundance and transcriptional activity in mesopelagic waters. Our findings provide the first physiological and genomic insights into the UBA868 group and suggest that members of this lineage contribute to the cycling of organic carbon, C1 compounds, and sulfur in marine ecosystems.

## Introduction

The vast majority of marine oligotrophic microorganisms remain uncultivated, limiting mechanistic understanding of their physiology and taxon-specific contributions to global biogeochemical cycles [1, 2]. Among these poorly characterized lineages, the order *Arenicellales* within *Gammaproteobacteria*, historically referred to as the UBA10353 marine group, represents a globally distributed yet still poorly understood bacterial lineage [3]. According to the latest Genome Taxonomy Database (GTDB; RS226), the order *Arenicellales* comprises thirteen families, among which *Arenicellaceae*, *Poriferihabitantaceae*, *Spongiicolonaceae*, and UBA868 represent the major lineages. These families occupy diverse ecological niches ranging from free-living planktonic environments (e.g., UBA868) to benthic habitats and symbiotic associations with marine eukaryotes (e.g., *Arenicellaceae* and *Poriferihabitantaceae*) [4]. Despite this broad ecological diversity, only three isolates have been cultured from the entire order to date, all belonging to *Arenicellaceae* [3, 5, 6], which remains the only validly published family in the order according to the List of Prokaryotic names with Standing in Nomenclature (LPSN; https://lpsn.dsmz.de/order/arenicellales). The scarcity of cultured representatives has therefore hindered experimental validation of genome-inferred metabolic traits within this lineage.

Environmental surveys have consistently highlighted the ecological importance of *Arenicellales*, particularly in oxygen minimum zones (OMZs). Members of this order were first reported as dominant bacterioplankton in the OMZs of the tropical Mexican Pacific [7] and were subsequently found to be enriched in depth-stratified waters of the Andaman Sea, Bay of Bengal, and Arabian Sea, where they often function as keystone taxa shaping microbial community structure [8]. Subsequent studies have revealed depth-dependent niche partitioning among *Arenicellales* members across marine water columns [4].

Within this order, the family UBA868 has recently gained attention due to its ecological significance but currently lacks cultured representatives. Sulfur oxidation represents an important energy-generating process in the mesopelagic ocean, yet the identities and physiological strategies of many key microbial players remain poorly resolved [9]. A recent study by Baltar et al. (2023) demonstrated a prominent role of UBA868 in sulfur and carbon cycling beneath the Ross Ice Shelf in Antarctica [10]. Their analyses further showed that UBA868 is a ubiquitous lineage that dominates the expression of key genes involved in sulfur oxidation and RuBisCO-mediated carbon fixation across the global mesopelagic ocean, suggesting a mixotrophic lifestyle coupling chemolithotrophy and heterotrophy. Similar metabolic strategies have also been reported in other globally distributed but underrepresented marine lineages, including SAR324, SUP05 (*Thioglobaceae*), and *Arcobacteraceae*, which are adapted to nutrient-limited marine environments through versatile mixotrophic lifestyles [11–13]. Together, these findings identify UBA868 as a potential contributor to global sulfur and carbon cycles and emphasize the need for cultured representatives to validate genome-inferred metabolic functions and ecological roles.

Here, we report the isolation of four strains representing a single novel species within the UBA868 family, including the first stably maintained culture of a heterotrophic representative, strain IMCC57338, from surface seawater of Garorim Bay, Yellow Sea. By integrating cultivation-based physiological characterization, genome-resolved metabolic reconstruction, and global environmental read recruitment, we provide mechanistic insights into the ecology and biogeochemical roles of this previously uncultured marine bacterial lineage.

## Materials and Methods

### Sampling and high-throughput cultivation

Seawater samples were collected at depths of 5 and 40 m from spatially distributed sites within Garorim Bay (37.01–37.03° N, 126.22–126.27° E), located in the Yellow Sea (West Sea of Korea), on four occasions between July 2023 and April 2024 (**Table S1**). Physicochemical parameters were measured *in situ* using a YSI 556 MPS multiprobe (YSI Inc., USA), and nutrient concentrations were determined from 0.45-µm filtered seawater samples using a HACH spectrophotometer (HACH Company, USA; **Table S1**).

High-throughput cultivation (HTC) based on the dilution-to-extinction approach was employed to isolate previously uncultured bacteria following established protocols [14, 15]. Briefly, seawater samples were diluted in low-nutrient heterotrophic medium (LNHM; **Table S2**) to ∼5 cells mL^−1^ and 1 mL aliquots were dispensed into 48-well microtiter plates (BD Falcon, USA) and incubated at 20. After four weeks, microbial growth was assessed by flow cytometry (Guava EasyCyte Plus, Millipore, USA) with SYBR Green I (Life Technologies, USA). Wells exhibiting growth (defined as cell densities >5.0×10^4^ cells mL^−1^) were cryopreserved in 10% (v/v) glycerol at −80 °C. LNHM was prepared from surface seawater (1 m depth) collected at the same site, filtered through a 0.22-μm membrane filter, autoclaved for 1.5 h, sparged with CO for 8 h, aerated for 24 h, and amended with nutrients (**Table S2**).

### Recovery and phylogenetic identification of UBA868 isolates

Phylogenetic identification of cultivated strains was based on nearly full-length 16S rRNA gene sequences amplified from cell lysates using InstaGene Matrix (Bio-Rad Laboratories, USA) with the primers 27F and 1492R. The resulting amplicons were Sanger sequenced with internal primers 518F and 800R (Macrogen Inc., Republic of Korea). Taxonomic classification was performed using the *classify.seqs* function in Mothur v1.39.5 [16] against the SILVA SSURef NR99 database (release 138.1) [17]. Of 3,252 bacterial strains recovered across all HTC experiments, four strains were affiliated with the family UBA868 within *Arenicellales* (UBA10353 marine group; **Table S3**). For 16S rRNA gene phylogeny, sequences affiliated with the UBA10353 marine group were retrieved from the SILVA database (release 138.1; **Table S4**), and a maximum-likelihood tree was inferred with RAxML v8.2.12 under the GTRGAMMA substitution model [18].

### Growth experiments on UBA868 representative IMCC57338

All four UBA868 isolates (IMCC57338, IMCC56312, IMCC55707, and IMCC58067) were revived from glycerol stocks in 20 mL of fresh LNHM and incubated under the same growth conditions as the initial HTC. Growth was monitored weekly for one month using Guava flow cytometry. Strain IMCC57338, which exhibited the most stable and reproducible growth, was selected as the representative strain for subsequent experiments. All growth assays for IMCC57338 were conducted in LNHM, and culture purity was routinely verified by 16S rRNA gene sequencing prior to each experiment.

Exponentially growing cultures of IMCC57338 (∼1.0×10^4^ cells mL^−1^) were inoculated into 20 mL of fresh LNHM for growth curve and temperature range experiments. Growth curves were determined at 20 °C with cell density measurements every three days. Specific growth rates (μ) were calculated by linear regression of ln-transformed cell densities during exponential phase, and doubling times were derived as ln(2)/μ. Temperature-dependent growth was assessed at 4–40 °C at weekly intervals. All experiments were performed in at least three independent biological replicates, and results are reported as mean ± SD.

### Morphological characterization by electron microscopy

Cellular morphology of IMCC57338 was examined by scanning electron microscopy (SEM) and transmission electron microscopy (TEM) following established protocols with minor modifications (**Supplementary methods**) [15, 19]. For SEM, cells from 1 L of exponential-phase culture (∼1.0×10^9^ cells) were collected on 0.22-µm polycarbonate membrane filters, fixed with glutaraldehyde, dehydrated through a graded ethanol series, chemically dried with hexamethyldisilazane (HMDS), carbon-coated, and imaged with a Hitachi S-4300 instrument (Japan). For TEM, exponential-phase cells were concentrated by centrifugation, stained with uranyl acetate, and imaged with a Talos L120C microscope (Thermo Fisher Scientific, USA).

### Genome sequencing, assembly, and functional annotation

Genomic DNA of IMCC57338 was extracted from centrifuged cell pellets using the DNeasy Blood & Tissue Kit (Qiagen, Germany). DNA from the three remaining UBA868 isolates was amplified from glycerol stocks by multiple displacement amplification (REPLI-g Single Cell Kit; Qiagen, Germany). Whole-genome libraries were prepared and sequenced by DNA Link, Inc. (Seoul, Republic of Korea) on the Illumina NovaSeq X Plus platform (2×150 bp paired-end; **Table S5**).

Raw reads were quality-filtered with BBDuk v39.01 to remove PhiX contamination, reads <100 nucleotides, and reads with Phred quality score <20. Filtered reads were assembled with SPAdes v3.15.5 using k-mer sizes of 21, 33, 55, 77, 99, and 127 [20]. Protein-coding sequences (CDSs) were predicted with Prodigal within the Distilled and Refined Annotation of Metabolism (DRAM) pipeline v1.4.6 [21], integrating KOfam, UniRef90, Pfam, CAZy, and MEROPS databases. KEGG Ortholog (KO) assignments were made with BlastKOALA [22] and KofamKOALA [23], and pathway completeness was determined with the KEGG-pathways-completeness tool (https://github.com/EBI-Metagenomics/kegg-pathways-completeness-tool) and KEGG Mapper. Pairwise average nucleotide identity (ANI) and average amino acid identity (AAI) were calculated with PyANI v0.3.0 and EzAAI v1.2.3, respectively [24, 25]. The metabolic map was created with BioRender (https://biorender.com).

### Phylogenomic analysis of the order *Arenicellales*

Reference genomes for phylogenomic analyses were compiled from three sources: 141 species-representative genomes classified as “o *Arenicellales*” from GTDB (release 226), 476 *Arenicellales*-affiliated MAGs from the OceanDNA catalog [10, 26], and 71 additional genomes retrieved from the Integrated Microbial Genomes (IMG). All genomes were de-replicated at a species-level threshold (ANI ≥95%) using dRep v3.5.0 [27]. Taxonomic assignments of the resulting non-redundant genomes were validated using GTDB-Tk v2.4.0 with the GTDB reference release R10-RS226. Genome quality was assessed with CheckM2 v1.0.1 [28], and only medium- to high-quality genomes (completeness >50%; contamination □10%) were retained (**Table S6**). Maximum-likelihood phylogenomic trees were inferred with RAxML v8.2.11 under the PROTGAMMAAUTO substitution model [18] and visualized using iTol v6.8.

### Metagenomic and metatranscriptomic read recruitment

The global distribution and transcriptional activity of the UBA868 group were assessed by recruiting reads from metagenomes (metaG) and metatranscriptomes (metaT) to the IMCC57338 genome (**Supplementary methods**). A total of 364 marine metagenomes from *Tara Oceans* [29], BioGEOTRACES [30], Malaspina [31], and two additional studies [32, 33] were analyzed (**Table S7**). Quality-filtered reads were mapped to the IMCC57338 genome at ≥95% ANI using CoverM v0.7.0 [34]. Ribosomal RNA gene regions were masked prior to mapping and relative abundances were normalized to total metagenomic read counts.

Transcriptional activity was evaluated using selected *Tara Oceans* metaT datasets [35] processed with the same quality-control and mapping framework as the metaG (**Table S8; Supplementary methods**). Ribosomal RNA reads were removed from each metaT dataset with RiboDetector v0.3. [36], and transcript abundances were expressed as reads per million (RPM). Differences in genomic and transcriptional abundance across pelagic depth zones were statistically evaluated with Wilcoxon rank-sum tests using the *ggpubr* package in R.

## Results and Discussion

### Cultivation and phylogenomic placement of UBA868 representatives

Application of dilution-to-extinction HTC [37] to Yellow Sea seawater collected in Garorim Bay yielded four bacterial isolates affiliated with the UBA10353 marine group (**Table S3**). The Yellow Sea has previously served as a productive source of oligotrophic marine bacteria, including the first cultured representatives of the SAR202 clade [15], underscoring the effectiveness of HTC for accessing uncultured bacterioplankton lineages.

High-quality genome sequences were obtained for all four isolates, each exhibiting >97% completeness and <1% contamination (CheckM2), with complete rRNA operons (5S, 16S, and 23S; **Table S5**). These genomes represent the most complete genomic resources currently available for members of the *Arenicellales* lineage (**Table S6**), with genome sizes of 2.64–2.70 Mbp, 2,526–2,619 predicted protein-coding genes, and consistent G+C content of 49.4–49.5%. A circular genome map of the representative strain IMCC57338 is shown in **Fig. S1**.

Phylogenomic analysis of 177 species-level reference genomes of *Arenicellales* (Table S6) confirmed placement of all four isolates within the previously uncultured family UBA868 (Fig. 1A, B), consistent with 16S rRNA gene phylogeny (Fig. S2) and prior classifications of *Arenicellales* as the UBA10353 marine group [4, 38]. All four isolates shared >99% 16S rRNA gene sequence identity and ANI values ≥97.5% (**Fig. 1C, D**), exceeding accepted bacterial species-delineation thresholds [39, 40], confirming that they represent a single novel species. Together, these results establish the first cultured bacterial species of the globally distributed UBA868 group.

**Figure 1.**
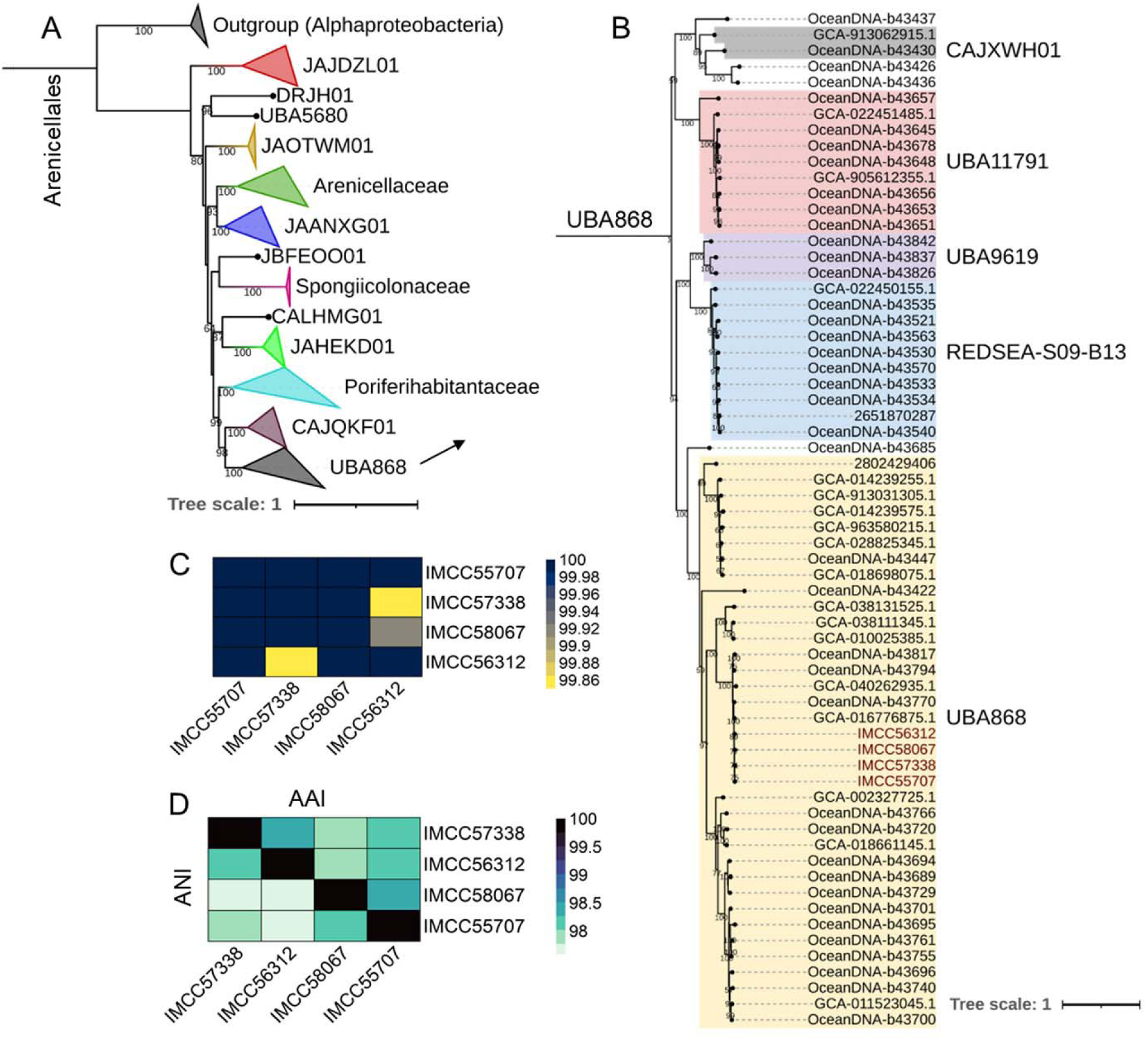
Phylogenetic placement and genomic relatedness of the four UBA868 isolates. (A) Maximum-likelihood phylogeny of major families within the order *Arenicellales* based on GTDB taxonomy (release 226). (B) Expanded phylogeny of the family UBA868 highlighting the four genomes recovered in this study (red font), with major genera indicated. Node names beginning with GCA, OceanDNA, and numeric-only identifiers, indicate genomes from the GTDB, OceanDNA MAG catalog, and IMG databases, respectively. Phylogenomic trees were inferred using RAxML v8.2.12 with the PROTGAMMAAUTO model from a concatenated alignment of 120 universal single-copy marker proteins generated by GTDB-Tk, with branch support assessed using 100 bootstrap replicates. Ten *Ruegeria* genomes (*Alphaproteobacteria*) were used as an outgroup (**Table S6**). Scale bar, one substitution per amino acid position. (**C**) Pairwise 16S rRNA gene sequence similarity (%) among the four UBA868 strains. (D) Pairwise average nucleotide identity (ANI; upper triangle) and average amino acid identity (AAI; lower triangle) values among the four genomes.

### Physiological and morphological characteristics of strain IMCC57338

Among the four isolates, strain IMCC57338 exhibited the most stable and reproducible growth and was selected for detailed physiological characterization. In LNHM, the bacterial culture entered stationary phase at day 18, reaching ∼2.05×10^6^ cells mL^−1^, followed by a decline after day 30 (**Fig. 2A**). The specific growth rate was 0.23 ± 0.01 day^−1^, corresponding to a doubling time of 2.93 ± 0.12 days, comparable to those of dominant marine oligotrophs, SAR11 subclade Ia (∼1.19–1.73 days) [41], SAR202 (∼2.88–3.85 days) [15], and SAR11 subclade V (AEGEAN-169; ∼1.45–1.93 days) [42], consistent with a slow-growing oligotrophic lifestyle.

**Figure 2.**
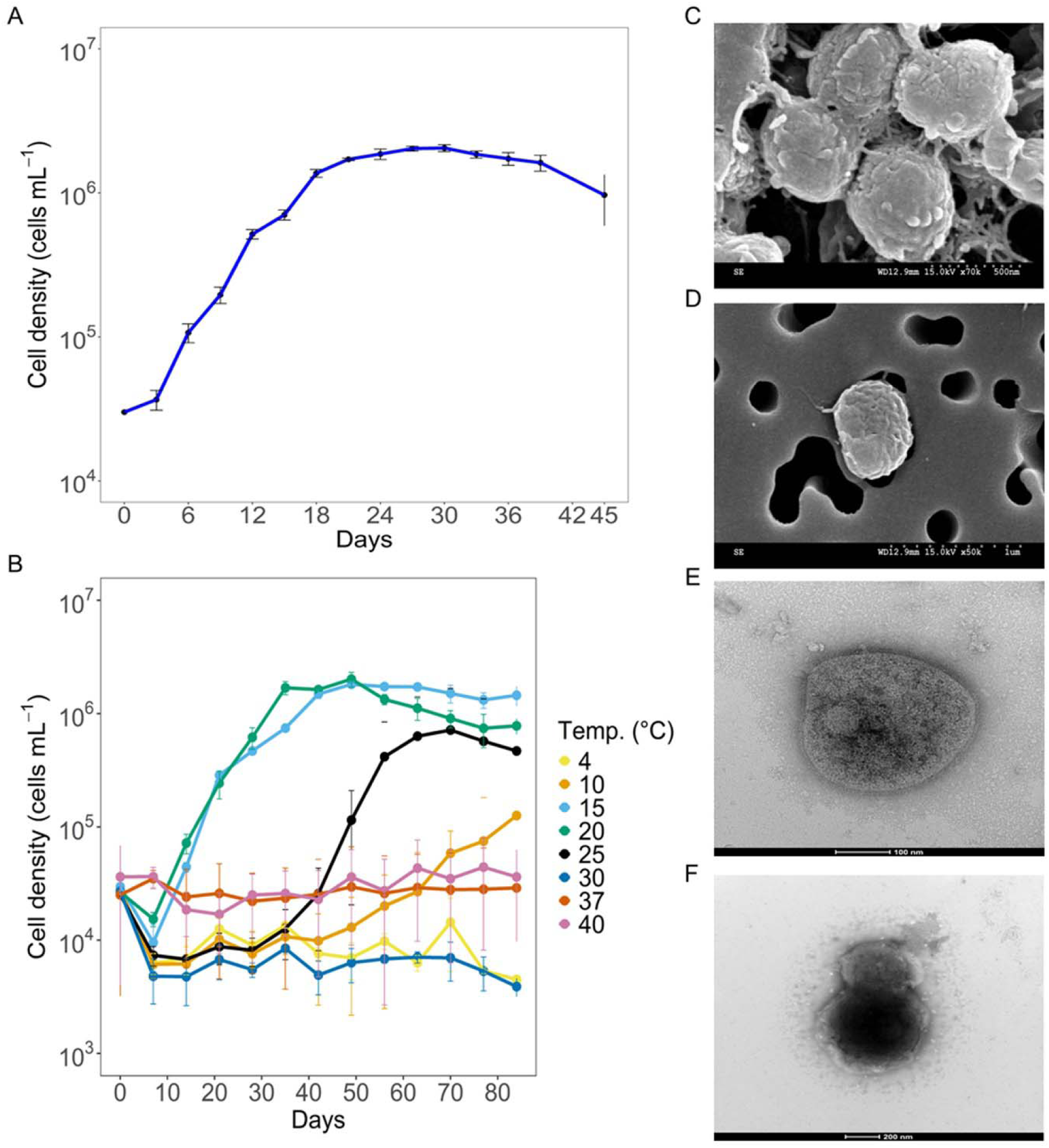
Growth and morphology of strain IMCC57338. (A) Growth curve in low-nutrient heterotrophic medium (LNHM) at 20 °C. (B) Temperature-dependent growth in LNHM across a range of 4–40 °C. Error bars represent the mean ± standard deviation (SD) from at least three independent experiments. (C, D) Scanning electron micrographs (SEM) showing coccoid cell morphology (scale bars, 0.5 µm and 1 µm, respectively). (E, F) Transmission electron micrographs (TEM) revealing cellular structure (scale bars, 100 nm and 200 nm, respectively).

Strain IMCC57338 exhibited optimal growth at 15–20 °C, whereas growth at 25 °C was preceded by a prolonged lag phase (**Fig. 2B**), suggesting that growth was near the upper temperature limit. SEM and TEM images revealed small coccoid cells with an average diameter of 0.47 ± 0.03 μm **(Figs. 2C–2F, S3**). No flagellar structures were observed, indicating a non-motile planktonic lifestyle.

### Central metabolism supports a chemoorganoheterotrophic lifestyle

Genome-based metabolic reconstruction of strain IMCC57338 (**Fig. 3**) showed highly conserved metabolic features across all four strains (**Table S9**). Central carbon metabolism encodes a nearly complete Embden–Meyerhof–Parnas (EMP) pathway, except for the absence of glucose kinase (*glk*; K00845 or *hk*; K00844, **Fig. 3**). No alternative glucose uptake or catabolism pathways, including the PtsG–Crr phosphotransferase system or the Entner–Doudoroff pathway, were detected in any genomes (**Table S9**). All genomes lacked isocitrate dehydrogenase (*icd*; K00031), yielding an incomplete tricarboxylic acid (TCA) cycle. Similar truncated “horseshoe-shaped” TCA configuration, also lacking α-ketoglutarate dehydrogenase, has been reported in *Candidatus Thioglobus autotrophicus* of the SUP05 clade [43] and is thought to favor biosynthetic carbon redirection over complete oxidation [44].

**Figure 3.**
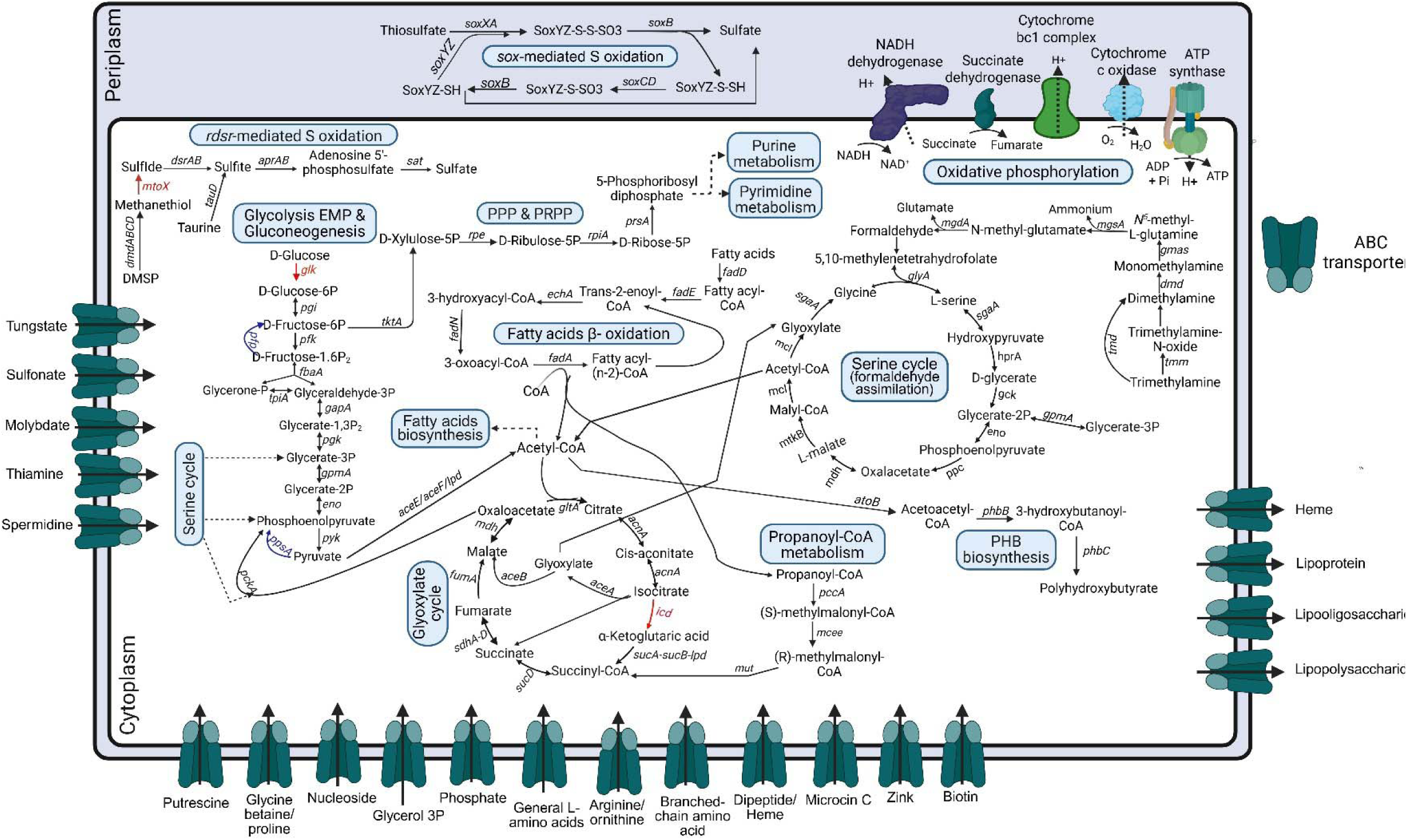
Metabolic reconstruction of strain IMCC57338. Schematic overview of major metabolic pathways inferred from genome annotation, emphasizing carbon, sulfur, and fatty acid metabolism. Gene names or steps shown in red are absent from the genome, although the remainder of the pathway is otherwise complete. Key gluconeogenetic steps are highlighted in blue. Expanded gene names, full pathway definitions, and completeness scores for each metabolic module are provided in **Tables S9** and **S11**. Created with BioRender.com.

In contrast, a complete glyoxylate shunt comprising isocitrate lyase (aceA; K01637) and malate synthase (aceB; K01638) was present in the genome, enabling bypass of TCA decarboxylation steps and growth on two-carbon compounds. The presence of a complete oxidative phosphorylation chain including cytochrome *c* oxidase supports aerobic respiration; the absence of high-affinity terminal oxidases (cbb_3_-type and bd-type) is consistent with adaptation to well-oxygenated environments [45]. No pathways for inorganic carbon or nitrogen fixation were detected, supporting a predominantly chemoorganoheterotrophic lifestyle. Although many UBA868 MAGs encode RuBisCO-based chemolithoautotrophic pathways [10], the cultivated strains described here are aerobic chemoorganoheterotrophs.

The complete absence of polysaccharide-degrading CAZymes, including glycoside hydrolases, polysaccharide lyases, and carbohydrate esterases (**Table S10**), further indicates a limited capacity for complex carbohydrate utilization, pointing to reliance on low-molecular-weight dissolved organic substrates [46, 47]. This is consistent with the presence of high-affinity ABC transporters for glycine betaine, amino acids, nucleosides, glycerol-3-phosphate, and small peptides (**Fig. 3**). In contrast, genes for fatty acid activation and β-oxidation support efficient fatty acid catabolism: even-chain fatty acids are fully degraded to acetyl-CoA, whereas odd-chain fatty acids additionally yield propionyl-CoA. Both intermediates are assimilated via the glyoxylate shunt and propionyl-CoA utilization pathways, which are fully encoded in IMCC57338. Similar reliance on fatty acid degradation has been reported in SAR86 members, including *Magnimaribacter mokuoloeensis* HIMB1674 [48]. The absence of flagellar biosynthesis genes is consistent with the observed non-motile microscopic morphology (**Fig. 2E, F**).

### Genomic evidence for methylotrophy and C1 assimilation

Metabolic reconstruction of IMCC57338 revealed a comprehensive suite of genes required for the oxidation of methylated amines (MAs), including trimethylamine *N*-oxide, trimethylamine, dimethylamine, and monomethylamine, supporting their utilization as one-carbon (C1) substrates (**Fig. 3**). These MAs, particularly trimethylamine *N*-oxide and trimethylamine, are widespread in marine environments at nanomolar to micromolar concentrations [49], and their utilization strategies vary across taxonomic lineages [50]. Rather than the canonical one-step methylamine dehydrogenase (*mau*) route, IMCC57338 employs an indirect pathway via γ-glutamylmethylamide and N-methylglutamate intermediates (Fig. 3), a strategy previously described in the methylotroph *Methylocella silvestris* [51]. The absence of methane monooxygenase (*mmo*) and methanol dehydrogenase (*mdh*) suggests that this metabolic capacity is specialized toward MAs rather than methane and methanol.

Beyond MA oxidation, IMCC57338 and the other three UBA868 genomes encode a complete serine cycle (**Table S9**), supporting potential for formaldehyde assimilation [52]. This pathway is likely replenished by glyoxylate derived from the glyoxylate shunt, with serine cycle intermediates funneling into gluconeogenesis to facilitate the biosynthesis of essential amino acids and nucleotides (**Fig. 3; Table S9**). Inclusion of methylamine in the HTC growth medium (**Table S2**) provides additional, albeit indirect, support for MA utilization in the culture. In contrast to dominant oligotrophic lineages such as SAR11 (*Pelagibacter ubique*) and SUP05 (*Thioglobus singularis*), which oxidize C1 substrates primarily for energy conservation [53, 54], IMCC57338 couples MA oxidation with formaldehyde assimilation, indicating a genomic capacity for “true” methylotrophy. Similar serine-cycle-mediated C1 assimilation has been tentatively identified in the uncultured SAR324 genomes [11], suggesting that this strategy may be more widespread among marine bacterioplankton than previously recognized.

### Sulfur metabolism in UBA868 heterotrophs

Reduced sulfur compounds serve as an important energy source for marine bacteria, particularly in oligotrophic regions where organic matter degradation continuously replenishes these sulfur species. Two complete inorganic sulfur oxidation pathways are encoded in the IMCC57338 genome (**Fig. 3**). The presence of the *soxXAYZBCD* gene cluster indicates the capacity of the strain to oxidize reduced sulfur compounds. This genomic evidence supports lithotrophic supplementation of heterotrophic carbon metabolism, a strategy that enhances carbon-use efficiency by reducing dependence on organic carbon for energy [55]. Notably, the presence of *soxCD* indicates direct oxidation to sulfate without sulfur globule accumulation, in contrast to *Thioglobus singularis* of the SUP05 clade, which lacks *soxCD* and consequently accumulates extracellular sulfur globules [19].

Strain IMCC57338 also encodes a complete reverse dissimilatory sulfite reductase (*rdsrAB*) pathway; the oxidative orientation of these DsrAB proteins was confirmed by their phylogenetic placement [56] (**Fig. S4**). The co-occurrence of both *sox*- and *rdsr*-mediated pathways in a single genome is uncommon but has been reported in other UBA868-affiliated MAGs [57] and certain *Roseobacter* clade members [58]. The presence of both pathways indicates the potential of UBA868 members to oxidize a range of reduced sulfur compounds, including elemental sulfur, thiosulfate, and sulfite, ultimately to sulfate [59]. Furthermore, IMCC57338 encodes genes for the degradation of organic sulfur compounds, including taurine (TauD/TfdA-family dioxygenase) and dimethylsulfoniopropionate (DMSP; DmdABCD demethylation pathway). These pathways likely generate reduced sulfur intermediates that channel into inorganic sulfur oxidation routes (**Fig. 3**), coupling organosulfur catabolism to energy conservation. While taurine cycling in the surface ocean is primarily attributed to SAR11 and *Roseobacter*, deep-ocean sulfur cycling is dominated by SAR324 and SUP05/Arctic96BD-19 lineages [60], suggesting that UBA868 may occupy an analogous niche in the mesopelagic ocean.

### Global prevalence and transcriptional dominance in the mesopelagic zone

The UBA868 group encompasses metabolic plasticity ranging from heterotrophic C1 and fatty acid utilizers to potential chemoautotrophs [10]. To assess the ecological relevance of the cultivated species, we performed read recruitment at ≥95% ANI to the IMCC57338 genome using metagenomic and metatranscriptomic data spanning major ocean basins and depth zones (**Fig. 4A**; **Table S7**).

**Figure 4.**
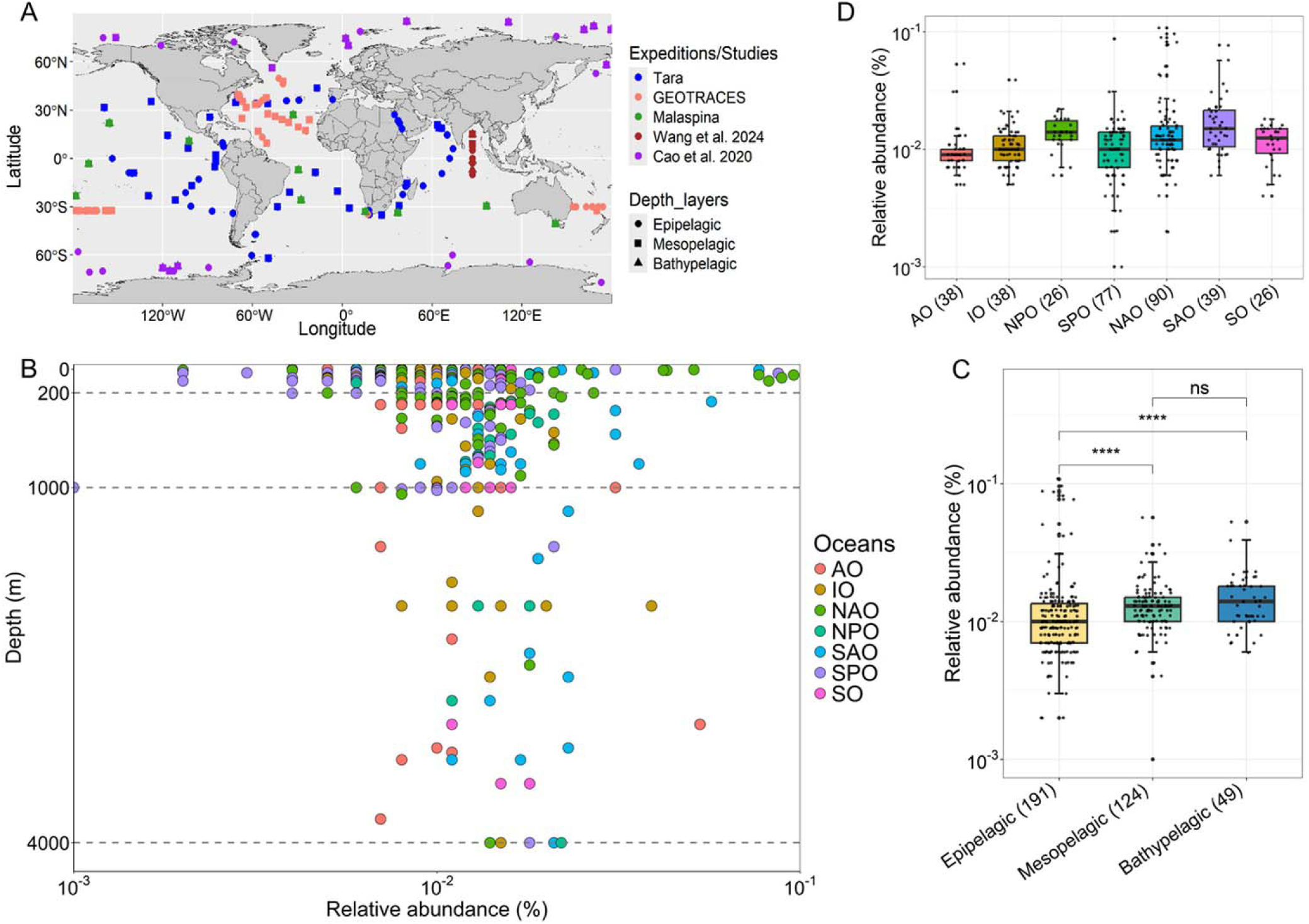
Global distribution and abundance of the IMCC57338 species inferred from metagenomic read recruitment. (A) Geographic distribution of metagenomic sampling stations from major oceanographic expeditions and studies, including *Tara Oceans*, GEOTRACES, and Malaspina [29–33]. (B) Vertical distribution of IMCC57338 relative abundance (%) plotted against sampling depth (m); points are colored by ocean basin (AO, Arctic Ocean; IO, Indian Ocean; NAO and SAO, North and South Atlantic Oceans; NPO and SPO, North and South Pacific Oceans; SO, Southern Ocean). (C) Comparative relative abundance across pelagic depth zones. Statistical significance was assessed using the Wilcoxon rank-sum test (ns, not significant; ****, *p*<0.0001). (D) Relative abundance across major ocean basins; sample numbers are indicated in parentheses. Metagenomic dataset details are provided in **Table S7**.

Although globally rare (mean relative abundance 0.014%), the IMCC57338 species was detected ubiquitously across ocean basins and depth zones, with local abundances exceeding 0.1% (**Fig. 4A–B; Table S7**), consistent with the broader UBA868 group distribution (global average ∼0.53%) [10]. Abundance of the species was consistently higher in aphotic waters, particularly the mesopelagic (200–1,000 m) and bathypelagic (1,000–4,000 m) zones, than in the epipelagic layer (0–200 m), where abundance was lower and more heterogeneous (pairwise Wilcoxon rank-sum tests, *p*≤0.001) (**Fig. 4B–C**). Geographic distributions were broadly comparable across basins, with consistent detection even in polar regions (**Fig. 4D**). Although IMCC57338 failed to grow below 15 °C in culture (**Fig. 2B**), sustained transcriptional activity of the UBA868 group beneath the Ross Ice Shelf [61] likely reflects intraspecific microdiversity, a common adaptive strategy for marine lineages spanning contrasting thermal regimes [62].

Metatranscriptomic read recruitment corroborated the observed abundance patterns, revealing a progressive increase in the transcriptional activity of the IMCC57338 species from surface waters down to the mesopelagic zone (**Fig. 5A–C; Table S8**). Transcript abundance was highest in the mesopelagic zone, followed by the deep chlorophyll maximum (DCM), with both layers showing significantly greater transcript levels than surface waters (*p*<0.01 and *p*<0.0001, respectively) (**Fig. 5D**). The mean mesopelagic transcript abundance (0.03%; **Table S8**) is comparable to the genomic abundance of SAR324 clade 2D (0.025%) in the same environment [63], underscoring the ecological prominence of UBA868 heterotrophs among midwater microbial assemblages. Paired metaT/metaG ratios were predominantly positive in the mesopelagic and negative in the epipelagic (**Table S8; Fig. 5E**), indicating disproportionately high transcriptional activity relative to genomic abundance in aphotic waters. Collectively, these results establish the UBA868 group as a metabolically active and ecologically relevant constituent of the mesopelagic microbial community.

**Figure 5.**
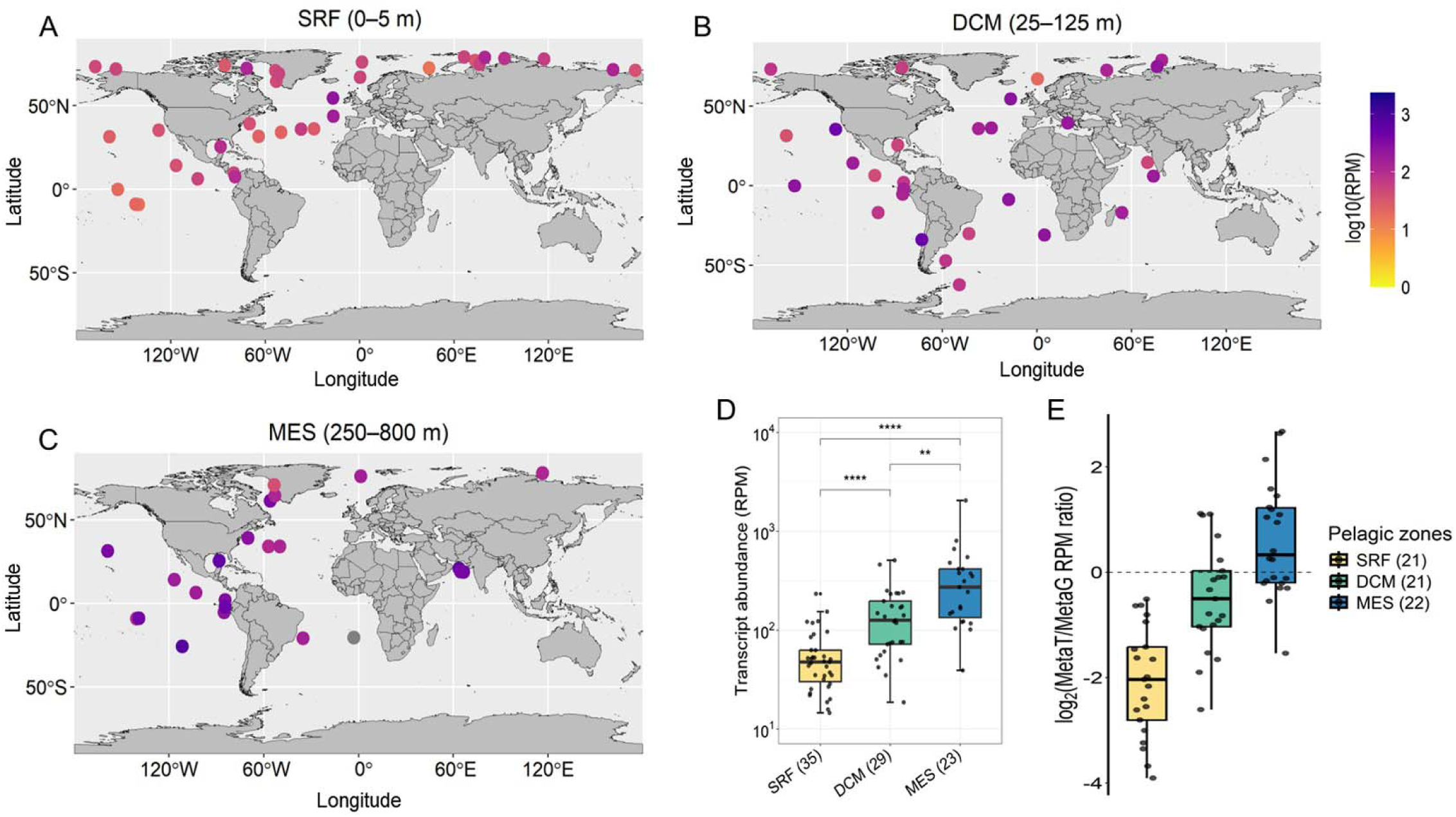
Global transcriptional activity of the IMCC57338 species inferred from metatranscriptomic read recruitment. (A–C) Metatranscriptomic read abundance (log_10_ RPM) mapped to the IMCC57338 genome for surface waters (SRF, A), deep chlorophyll maximum (DCM, B), and the mesopelagic zone (MES, C) sampled during the *Tara Oceans* expedition. Each point represents an individual sampling station; color scale indicates transcript abundance (reads per million; RPM). (D) Comparative transcript abundance (RPM) by pelagic zones. (E) Log_2_(metaT/metaG RPM) ratios for samples with paired metatranscriptomic and metagenomic datasets, illustrating transcriptional activity relative to genomic abundance across depth layers. Statistical significance was assessed using the Wilcoxon rank-sum test: ns, not significant; **, *p* < 0.01; ****, *p* < 0.0001. Metagenomic and metatranscriptomic dataset details are provided in **Table S8**.

### Conclusion and taxonomic proposal

Our study provides the first integrated genomic, physiological, and ecological characterization of heterotrophic members of the UBA868 group. The cultivated strain IMCC57338 reveals hallmark oligotrophic adaptations, including a streamlined genome, slow growth, and reliance on low-molecular-weight substrates, combined with metabolic versatility encompassing methylotrophy and sulfur-based lithoheterotrophy. Global read recruitment confirms that this species is widely distributed and transcriptionally dominant in the mesopelagic ocean. Future work should prioritize expanded cultivation targeting potential autotrophic UBA868 members and experimental validation of sulfur oxidation and methylotrophic activities.

Based on the distinct phylogenetic position of the UBA868 group (equivalent to GTDB g UBA868) and its demonstrated ecological relevance, we propose the name *Mediimaricoccus* gen. nov. for this lineage, with *Mediimaricoccus garorimensis* sp. nov. as the type species, and *Mediimaricoccaceae* fam. nov. with *Mediimaricoccus* as the type genus. Formal taxonomic descriptions and protologues are provided in the **Supplementary Information**.

## Data availability statement

Raw sequences and genome assemblies generated in this study have been deposited in the NCBI SRA and GenBank under BioProject PRJNA1395325. GenBank accession numbers are JBUBJZ000000000 (IMCC57338), JBUBKA000000000 (IMCC55707), JBUBKB000000000 (IMCC58067), and JBUBKC000000000 (IMCC56312). Strain IMCC57338 is available from the Cho Laboratory (https://www.cholabinha.org/) upon request.

## Acknowledgements

We thank Prof. Aharon Oren (The Hebrew University of Jerusalem, Israel) for advice on taxonomic nomenclature.

