## Supplementary Information for "Cultivation and genomic characterization of the first representative of the globally distributed marine UBA868 group"

**Rajeev *et al.***

**This file includes:**

**Supplementary Methods……………………………………………………Pages 2–3**

**Supplementary Taxonomic Descriptions……………………………… Pages 4–5**

**Supplementary Figures……………………………………………………… Pages 6–9**

**Supplementary References…………………………………………………. Page 10–11**

### Supplementary Methods

#### Scanning and transmission electron microscopy

Scanning electron microscopy (SEM) was performed following a previously described protocol for heterotrophic members of the SUP05 clade [1], with minor modifications. Cells of strain IMCC57338 were harvested from 1 L of exponentially growing culture (cell density ~$1.0 \times 10^6$ cells $mL^{-1}$; total ~$1.0 \times 10^9$ cells) and gently filtered onto a 0.22-μm pore size polycarbonate membrane filter (Millipore) under low vacuum using a peristaltic pump. Immediately after filtration, 1 mL of 0.1 M sodium cacodylate buffer containing 2% (v/v) glutaraldehyde was applied to the filter and incubated for 1 h at 4 °C to fix cellular ultrastructure. Following fixation, cells were dehydrated through a graded ethanol series (30%, 50%, 70%, 90%, and twice in 100% ethanol), with each step incubated for 10 min prior to gentle solvent removal. Chemical drying was then performed using sequential combinations of absolute ethanol and hexamethyldisilazane (HMDS; Sigma-Aldrich) at ethanol:HMDS ratios of 2:1, 1:1, and 1:2, followed by a final incubation in 100% HMDS. After complete drying, the filter was mounted onto an aluminum SEM stub, carbon-coated, and examined using a scanning electron microscope (SEM; S-4300; Hitachi, Japan) for high-resolution imaging of cell morphology.

For transmission electron microscopy (TEM), exponentially growing cells of strain IMCC57338 were harvested from 1 L of LNHM culture (total cell number ~$1.0 \times 10^9$) by centrifugation at 12,000 rpm for 90 min at 4 °C using a Sorvall RC 6 Plus superspeed centrifuge (Thermo Scientific, USA). The resulting cell pellet was gently resuspended in 2 mL of sterile LNHM. A small aliquot of the concentrated cell suspension was applied to Formvar/carbon-coated copper grids and allowed to adsorb for 1–2 min. Excess liquid was removed by gentle blotting with filter paper, and the grids were immediately negatively stained with freshly prepared 2% (w/v) uranyl acetate for 1–2 min in the dark. After staining, excess uranyl acetate was blotted off and the grids were air-dried at room temperature. Electron micrographs were acquired at an accelerating voltage of 120 kV using a transmission electron microscope (Talos L120C, Thermo Fisher Scientific, Waltham, USA) facilities available at National Instrumentation Center for Environmental Management (NICEM), Seoul, Republic of Korea.

### Recruitment of metagenomic and metatranscriptomic reads

The global distribution and transcriptional activity of the UBA868 group were inferred by recruiting metagenomic (metaG) and metatranscriptomic (metaT) reads to the genome of the cultivated representative strain IMCC57338. In total, 364 publicly available MetaG datasets were analyzed, including those from previous studies (n = 85) [2, 3] and from major oceanographic expeditions such as *Tara Oceans* (prokaryote-enriched; n = 127) [4], BioGEOTRACES (n = 98) [5], and Malaspina (n = 54) [6] as detailed in **Table S7**. In cases where multiple sequencing replicates were available (e.g., *Tara Oceans* and Malaspina datasets), the replicate with the highest sequencing depth was selected. These metagenomes collectively span diverse ocean basins and depth zones, providing broad representation of UBA868 distribution.

All metaG underwent quality control using BBduk v39 [7], including adapter trimming, contaminant removal, exclusion of low-quality bases (Phred < 20) and short reads (< 100 bp), and removal of phiX174 sequences, following our previously described protocol [8]. Quality-filtered reads were mapped to the IMCC57338 genome using CoverM v0.7.0 with the following parameters: --min-read-aligned-length 50, --min-read-percent-identity 95, --min-read-aligned-percent 60, and --min-covered-fraction 0 [9]. Ribosomal gene regions in the reference genome were masked prior to mapping to prevent recruitment to conserved regions. Relative abundance was computed using the relative_abundance method in CoverM, normalizing to total metagenomic reads to estimate the proportion of reads recruited to isolate IMCC57338 within each metagenome.

Transcriptional activity of cultured representative strain IMCC57338 was assessed by mapping metaT reads from *Tara Oceans* [10]. Selected metaT datasets (**Table S8)** were processed using identical quality control, mapping parameters and tools as metaG, except that reads ≤ 50 bp were discarded and rRNA sequences were removed with RiboDetector v0.3.1 [11] before mapping. The resulting mapped reads were normalized and expressed as reads per million metatranscriptomics reads (RPM) to enable quantitative comparison across samples.

### Supplementary Taxonomic Descriptions

#### Proposal of ranks of the new taxa

#### Description of *Mediimaricoccus* gen. nov.

*Mediimaricoccus* (Me.di.i.ma.ri.coc'cus. L. masc. adj. *medius*, middle; L. neut. n. *mare*, sea; N.L. masc. n. *coccus* (from Gr. masc. n. *kokkos*, a grain, a seed), a coccus, a spherical bacterium; N.L. masc. n. *Mediimaricoccus*, a marine spherical bacterium associated with mid-water oceanic environments).

Cells are coccoid, small, non-motile, and occur as free-living planktonic cells. Members of the genus are aerobic, oligotrophic, and chemoheterotrophic. Growth occurs in seawater-based liquid media under low-nutrient conditions. Cells exhibit slow growth typical of marine oligotrophic bacteria.

The genus *Mediimaricoccus* represents a phylogenetically distinct lineage of marine *Gammaproteobacteria* based on whole-genome phylogenomics and is equivalent to GTDB g__UBA868. The type species of the genus is *Mediimaricoccus garorimensis.*

#### Description of *Mediimaricoccus garorimensis* sp. nov.

*Mediimaricoccus garorimensis* (ga.ro.ri.men'sis. N.L. masc. adj. *garorimensis*, pertaining to Garorim Bay, Yellow Sea, Republic of Korea, from where the type strain was isolated).

In addition to the characteristics described for the genus *Mediimaricoccus*, the species is described as follows. Cells are uniformly coccoid, with no rod-shaped or elongated morphotypes observed. Cells occur singly or in pairs. The organism grows under aerobic conditions in seawater-based oligotrophic media. Optimal growth occurs at temperatures between 15 and 20 °C.

The type strain, IMCC57338$^{T}$, was isolated from surface seawater of Garorim Bay, West Sea, Republic of Korea, at a depth of 5 m. The genome size of the type strain is 2.75 Mbp, with a DNA G + C content of 49.2 mol%, as determined from the genome sequence. The GenBank accession number for the genome sequence of strain IMCC57338$^{T}$ is JBUBJZ000000000.

**Description of *Mediimaricoccaceae* fam. nov.**

*Mediimaricoccaceae* (Me.di.i.ma.ri.coc.ca.ce'ae. N.L. masc. n. *Mediimaricoccus*, a bacterial genus name; -*aceae*, ending to denote a family; N.L. fem. pl. n. *Mediimaricoccaceae*, the *Mediimaricoccus* family).

The description of the family *Mediimaricoccaceae* is the same as that of the genus *Mediimaricoccus*. The type genus is *Mediimaricoccus*. Members of this family belong to the class *Gammaproteobacteria* and are assigned to the order *Arenicellales* (UBA10353 marine group) based on 16S rRNA gene phylogeny and whole-genome phylogenomic analyses. The family corresponds to GTDB f__UBA868.

### Supplementary Figures

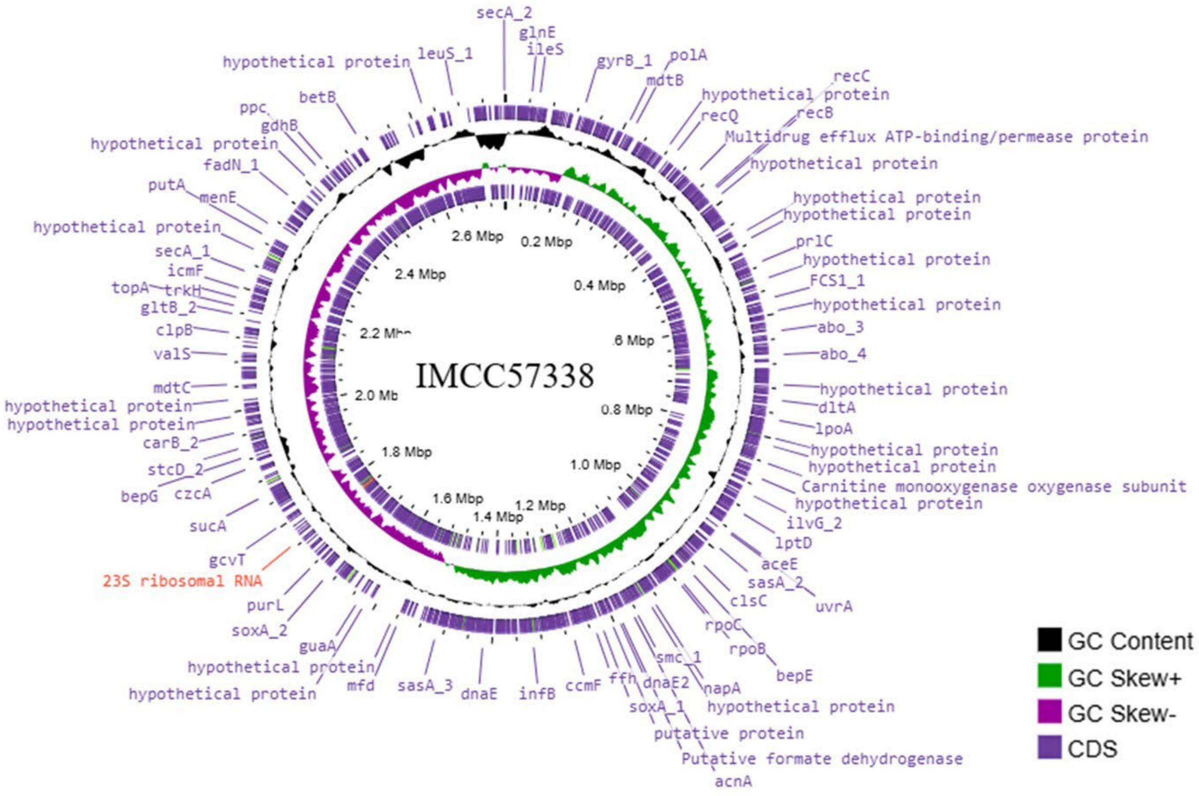


**Supplementary Figure S1.** Genome map of strain IMCC57338. Circular genome map of the UBA868 cultured representative strain IMCC57338. The genome map was visualized using online Proksee tool based on Prokka genome annotation. From the outermost ring to the center, the map displays predicted coding sequences (CDSs), GC content, GC skew, and the genome size scale.

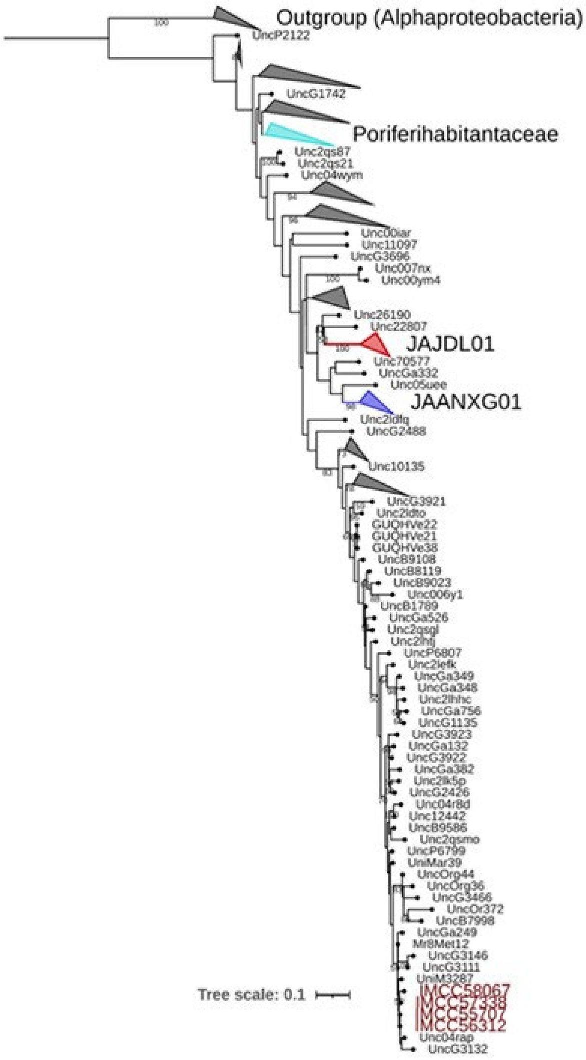


**Supplementary Figure S2.** 16S rRNA gene–based phylogenetic placement of UBA868 strains obtained in this study. Maximum-likelihood phylogenetic tree showing the relationships of the four strains recovered in this study (marked in red) based on 16S rRNA gene sequences. Reference sequences affiliated with the order UBA10353 marine groups were retrieved from the SILVA SSURef NR99 database (release 138.1). The phylogeny was inferred using RAxML v8.2.12 under the GTRGAMMA substitution model, with branch support assessed using 100 bootstrap replicates. Colored branches indicate corresponding GTDB taxonomy based on SBDI Sativa curated 16S GTDB database (https://github.com/biodiversitydata-se/sbdi-gtdb). Ten 16S rRNA gene sequences from the genus *Ruegeria* (Alphaproteobacteria) were used as an outgroup to root the tree. Scale bar indicates 0.1 substitutions per nucleotide position.

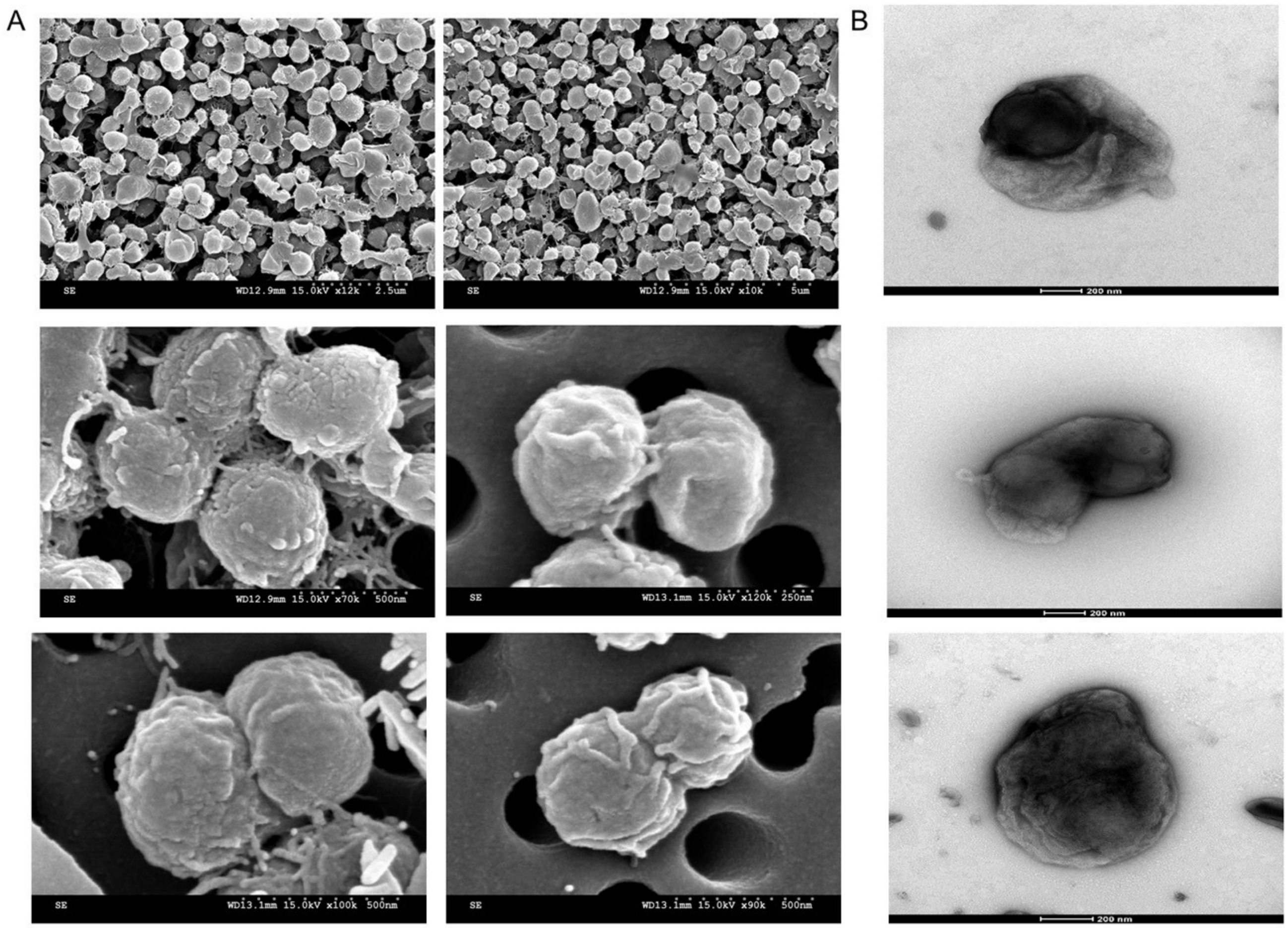


**Supplementary Figure S3.** Microscopic visualization of UBA868 heterotrophic strain IMCC57338. (A) Scanning electron micrographs (SEM) showing the morphology of IMCC57338 cells. (B) Transmission electron micrographs (TEM) revealing the cellular ultrastructure of IMCC57338.

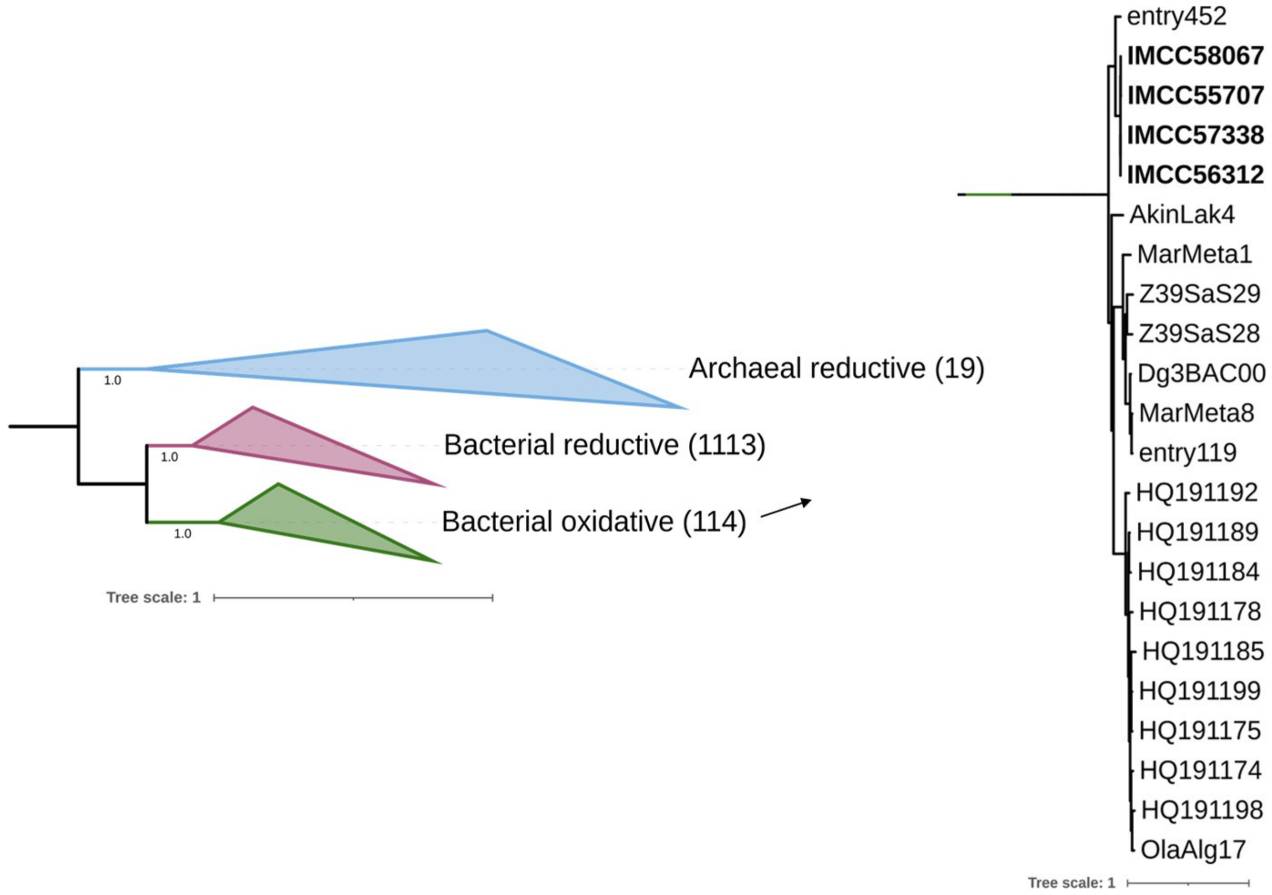


**Supplementary Figure S4.** Phylogenetic positioning of reverse/oxidative dissimilatory sulfite reductase (rDsrAB) sequences identified in the four UBA868 genomes. Reference DsrAB protein sequences representing reductive and oxidative natures were retrieved from Pelikan et al. (2016) [12] and aligned together with UBA868 rDsrAB sequences using MAFFT. The IMCC strains are shown in bold. The maximum-likelihood phylogenetic tree was inferred using FastTree v2.1.11 with the LG the LG + Γ (gamma) model. Tree visualization and annotation were performed using iTOL v6.8.
